# Rigid geometry solves “curse of dimensionality” effects in clustering methods: An application to omics data

**DOI:** 10.1101/094391

**Authors:** Shun Adachi

## Abstract

The quality of samples preserved long term at ultralow temperatures has not been adequately studied. To improve our understanding, we need a strategy to analyze protein degradation and metabolism at subfreezing temperatures. To do this, we obtained liquid chromatography-mass spectrometry (LC/MS) data of calculated protein signal intensities in HEK-293 cells. Our first attempt at directly clustering the values failed, most likely due to the so-called “curse of dimensionality”. The clusters were not reproducible, and the outputs differed with different methods. By utilizing rigid geometry with a prime ideal *I*-adic (*p*-adic) metric, however, we rearranged the sample clusters into a meaningful and reproducible order, and the results were the same with each of the different clustering methods tested. Furthermore, we have also succeeded in application of this method to expression array data in similar situations. Thus, we eliminated the “curse of dimensionality” from the data set, at least in clustering methods. It is possible that our approach determines a characteristic value of systems that follow a Boltzmann distribution.

## Introduction

Even when frozen, biological samples degrade during aging, and most frozen cell cultures are stored for only two years. However, the cause of degradation is not well understood. A few reports have described enzymatic activities in frozen cultures, including lipase and peroxidase activities (see, e.g., [1, 2]). However, the proteomic details of cells stored at subfreezing temperatures are not known. One study used liquid chromatography-mass spectrometry (LC/MS) to study frogs in a simulated winter environment [3]; it lacks solid statistical analysis and did not consider subfreezing temperatures. In order to evaluate the potential degradation/metabolism, it is important to obtain solid proteomic data from actual frozen cultures under long-term storage in a subfreezing environment.

To do this, we devised a procedure that can distinguish samples from long-term storage from those that have been freshly prepared. Clustering analysis is a popular approach to such an evaluation; this approach uses criteria for similarity/dissimilarity to divide the data into meaningful groups. It is based on a bottom-up calculation of the data, and thus the criteria are part of the system. However, it is still necessary to define the groups and to select the actual clustering methods. If different clustering methods yield the same topological structure of the hierarchical tree or the same indices of the clusters, it can be assumed that the output of the analysis is sound; however, this is not always achieved, and discrepancies cast doubt on the results.

There are two types of clustering analysis: hierarchical clustering [4] and nonhierarchical clustering [5]. Hierarchical clustering is appropriate when there is distance/dissimilarity between the data points; it joins data based on similarities to a given point, and then combines those points based on their similarities. In this way, a multidimensional data set is reduced to a two-dimensional set, with the axes indicating labeling and clustering distance. Representatives of these methods include simple linkage, complete linkage, group averaging, weighted averaging, methods using the centroid or median, and Ward’s method. If we define the dissimilarity of *i, j,* and *k* to be *C_i_, C_j_*, and *C_k_*, respectively, then

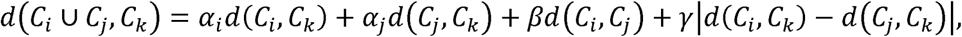

where the values of α_*i*_, α_j_, β, and □ are given in Table 1.

**Table 1.** Parameters of various hierarchical clustering methods. Β, □: corrections based on the triangle *ijk*; Dissimilarity [*E2:* Euclidean distance; *E2’*: half of the squared Euclidean distance]; Monotony [monotonically increasing lengths (We note that this is not true in the centroid and median methods. The value of a monotonic increase depends on the particular situation.); T: true; F: false;]; Metric [expansion & reduction: renewal of ongoing clustering by increasing or reducing the distance between data points].

| method | $\alpha_i$ | $\alpha_j$ | $\beta$ | □ | Dissimilarity | Monotony | Metric |
| --- | --- | --- | --- | --- | --- | --- | --- |
| single | 1/2 | 1/2 | 0 | -1/2 | no<br>restriction | T | reduction |
| complete | 1/2 | 1/2 | 0 | 1/2 | no<br>restriction | T | expansion |
| group<br>average | $\frac{n_i}{n_i + n_j}$ | $\frac{n_j}{n_i + n_j}$ | 0 | 0 | no<br>restriction | T | conserved |
| weighted<br>average | 1/2 | 1/2 | 0 | 0 | no<br>restriction | T | conserved |
| centroid | $\frac{n_i}{n_i + n_j}$ | $\frac{n_j}{n_i + n_j}$ | $\frac{-n_i n_j}{(n_i + n_j)^2}$ | 0 | $E2$ | F | conserved<br>&<br>reduction |
| median | 1/2 | 1/2 | -1/4 | 0 | $E2$ | F | conserved<br>&<br>reduction |
| Ward | $\frac{n_i + n_k}{n_i + n_j + n_k}$ | $\frac{n_j + n_k}{n_i + n_j + n_k}$ | $\frac{-n_k}{n_i + n_j + n_k}$ | 0 | $E2'$ | T | conserved<br>&<br>expansion |

Nonhierarchical clustering, for example, the *k*-means method, is an optimization approach based on classification. The number of groups, *k*, is initialized. Dissimilarity is measured using the squared Euclidean distance. The *k* groups are then determined and scored as each data point is added. The grouping that gives the lowest score is chosen, and the process is repeated. The selection of the number of groups is a top-down approach, but from other aspects, this is a bottom-up approach. Each of the eight methods, including seven hierarchical methods and the *k*-means method, can be relatively easily implemented on a computer, and these are more frequently used than are other, more complex methodologies.

A problem arises, however, when there is a high-dimensional data set (more than 1000 variables); this effect is referred to as the “curse of dimensionality”. In such a situation, the variances among samples become large and sparse, and a clustering analysis produces meaningless results (see, e.g., [6]). One solution is to use machine learning [7]. However, the high dimensionality produces a very large or incalculable value for the Akaike information criterion, and this means the solution may be invalid. Alternatively, principal component analysis (PCA) with maximized variances of variance-covariance matrix [8] or non-metric multi-dimensional scaling calculating similarity in reduced dimensions without the constraint of linearity (nMDS) [9] is utilized to classify the observed data sets. These methods also exhibited “curse of dimensionality” in very high dimension, however.

In this manuscript, we present a mathematical solution that uses rigid geometry to pretreat the data prior to a cluster analysis. The most important aspect of clustering is determining the metric, and we considered the p-adic metric (of a prime ideal), which is based on rigid geometry (Please refer S1 Appendix). Utilizing the idea of blowing up singularity in rigid geometry, one can resolve singularity that affects most parts of fluctuations and purify the internal characteristic residing in the dataset [10]. This allowed us to discriminate between control samples and those that had been held in long-term storage. An appropriate metric must satisfy (i) a separation axiom (not necessarily nonnegative), (ii) the identity of indiscernibles, (iii) symmetry, and (iv) the triangle inequality. Examples of metrics include the absolute distance, the Chebyshev distance, the Euclidean distance, the average Euclidean distance, squared Euclidean distance, the Minkowski distance, the correlation efficiency, cosine efficiency nearest neighbor distance and the radial basis kernels. The selection of the metric significantly affects the calculation (see, e.g., [6]). We picked up nearest neighbor distances 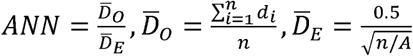 and radial basis kernals 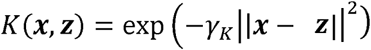 together with Euclidean distances and correlation distances *d_cor_(**x,z**)* = 1-*r_cor_* (***x***, ***z***)[11] to compare the results with a *p*-adic metric we invented, when *d_i_* is an element *i* of distance vector, *A* is a total study area, ***x**, **z*** are a vectors of interests and *r_cor_* is the correlation of them. You can easily see that all the nearest neighbor distances, radial basis kernels and Euclidean distances preserve the high dimensionalities in the distance vector ***d**.* Correlation distances seems to cancel out high dimensionality with quotient, but still does not erase the trace of singularities. This might be the cause of “curse of dimensionality”. A principle of modern geometry is that there exists a nilpotent state in which observed values converge and oscillate about -1; this results in a system that is easier to handle. Rigid geometry is a well-known mathematical field and is based on the complete non-Archimedean field; it was introduced in 1962 by John Tate [12–14]. It allows us to use *p*-adic elliptic curves to solve the singularity problem [10]. In this system, the non-Archimedean valuation system enables values to converge globally, but locally the values are free, due to the high dimensionality. There is a mathematically well-known topology that meets this type of requirement, Grothendieck topology (G-topology) [15]. Below, we will present the application of rigid geometry to a biological data set, and we will show that this removes the effects of the “curse of dimensionality”; the resulting topology of the clusters can then be easily interpreted.

## Results

Direct analysis of unused LC/MS data resulted in nonproper clustering of samples with clustering methods or non-metric multidimensional scaling We extracted proteins from HEK-293 cells that had been stored in different freezing conditions. As a control, we collected fresh samples from a cell culture, samples from cells frozen for 1 h at -80°C (1h), and samples from cells frozen overnight at -80°C and subsequently transferred to liquid nitrogen and held overnight (o/n-o/n). For the treated samples, we used samples that had been preserved in liquid nitrogen for 2 or 3 years. See the Methods section for more details. After performing LC/MS, we extracted unused values (the amount of total, unique peptide evidence related to a given protein) and performed clustering analyses using various hierarchical methods and the *k*-means method. We found that although different treatments of the sample gave significantly different results, clustering did not yield meaningful information; the clusters were not reproducible, and the outputs differed with different methods; see Fig 1A. The control and treatment data were combined and neural-network-based machine learning produced clustering of cl values (the expected number of columns in the model of the training data set) on both the control samples and the storage samples. The samples that had been stored for 2 years were distinct from the control values, and only those stored for 3 years were interspersed (Fig 1A). This suggests the presence of “curse of dimensionality” effects (see, e.g., [6]). Additionally, PCA exhibited 89% of the data can be explained by a single component (PC1) with absolute correlation values > 0.74 in each, indicating failure in clustering (Table 2). PC2 and PC3 occupies 6% and 1% contributions, respectively. nMDS did not show proper clustering among control samples and samples frozen for years, indicating failure (Fig 1A). The number of unknown parameters in the neural network was 1636. This idea is illustrated in Fig 2A, and actual structures in geometric space are important for conducting these calculations [6]. In this first analysis, we used the simple unused values with Euclidean distances when calculating the dissimilarity.

**Fig 1.**
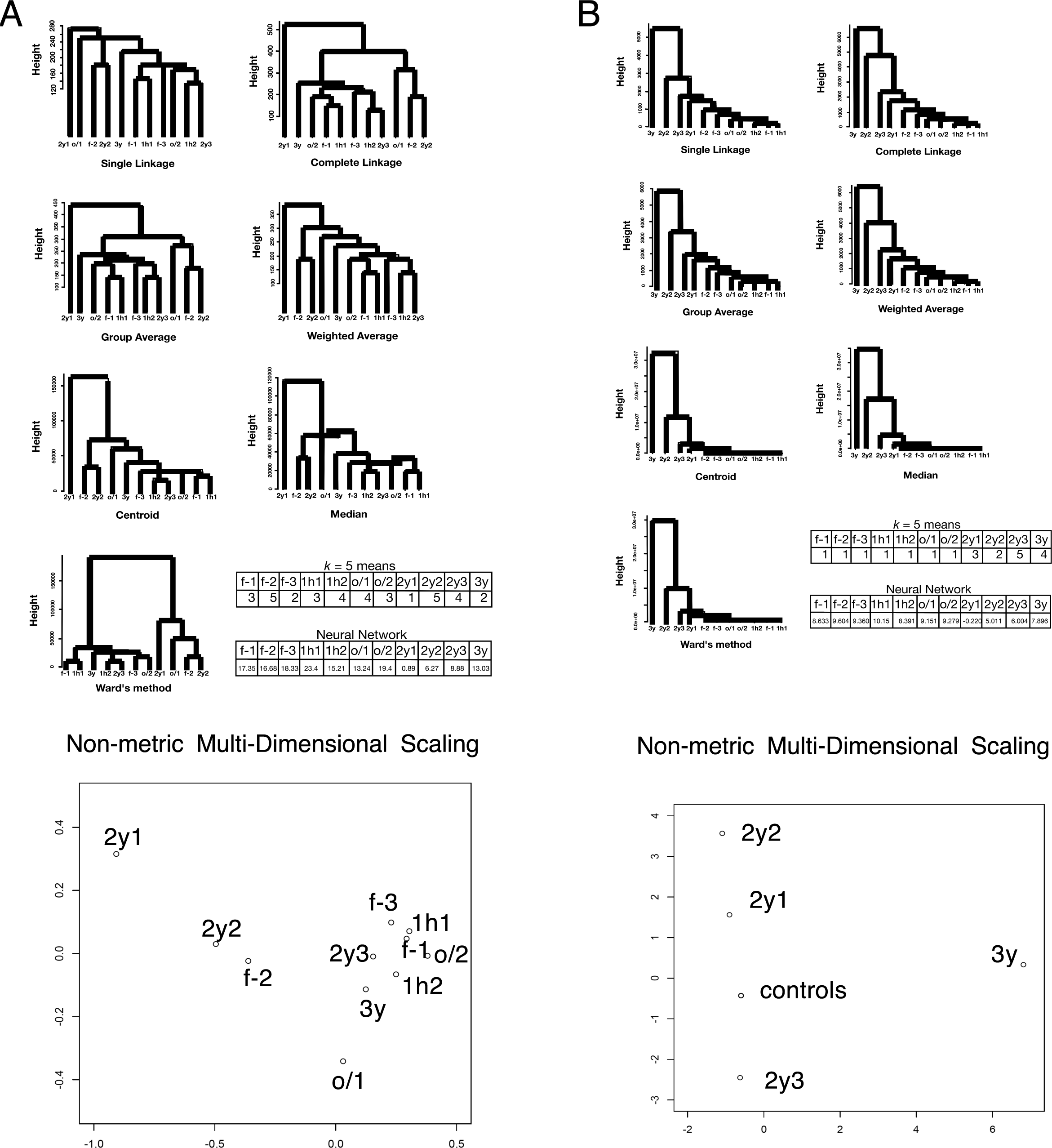
Clustering of value sets for each protein in the LC/MS data of HEK-293 (*N*= 803). f-1, f-2, and f-3: freshly prepared samples in replicates 1, 2, and 3, respectively; 1h1 and 1h2: samples frozen at -80°C for 1 h in replicates 1 and 2, respectively; o/1 and o/2: samples that remained at -80°C overnight and then in liquid nitrogen overnight (“o/n-o/n”) in replicates 1 and 2, respectively; 2y1, 2y2, and 2y3: samples preserved in liquid nitrogen storage according to the RIKEN protocol for approximately 2 years in replicates 1, 2, and 3, respectively; 3y: sample preserved in liquid nitrogen storage according to the RIKEN protocol for approximately three years. The numbers in the *k*-means table are the indices of the classified groups. The numbers in the neural network table are the cl factors, which represents the one-dimensional characteristics of the systems. The differences in cl values show dissimilarity of the samples. Please also see the Methods section. (A) Unused values. (B) *v* values.

**Table 2.** Correlation matrix of LC/MS data and principal components.

| raw | f-1 | f-2 | f-3 | 1h1 | 1h2 | o/1 | o/2 | 2y1 | 2y2 | 2y3 | 3y |
| --- | --- | --- | --- | --- | --- | --- | --- | --- | --- | --- | --- |
| PC1 | 0.97 | 0.92 | 0.97 | 0.97 | 0.98 | 0.95 | 0.96 | 0.74 | 0.88 | 0.98 | 0.96 |
| PC2 | -0.11 | 0.28 | -0.10 | -0.12 | -0.12 | -0.08 | -0.20 | 0.64 | 0.40 | -0.00 | -0.12 |
| PC3 | -0.02 | 0.19 | -0.12 | -0.00 | -0.01 | 0.23 | -0.07 | 0.17 | 0.13 | -0.00 | -0.12 |

## A *p*-adic metric based on rigid geometry eliminated “curse of dimensionality” effects with LC/MS data

To avoid the pitfalls described above, we designed a better metric for the calculation of groups (Please refer S1 Appendix). If we choose an appropriate geometric metric that is nilpotent for convergence/divergence of values and converges to an oscillation around -1, then more over-converged output can be extracted and used to discriminate between the observed characteristics. A common approach for this is to use rigid geometry. When using a *p*-adic metric that includes a subring of norm < |1|, a non-Archimedean field is more likely to converge than is an Archimedean real field or a complex field. The geometry converges globally, but locally the values are free, enabling freedom from the restriction due to “curse of dimensionality”.

One example of this type of analysis is illustrated in Fig 2B (see, e.g., [16]). Consider characteristic points of an icosahedron projected onto a sphere: 12 vertices (indicated in blue in Fig 2B), 20 barycenters (the 20 centers of the triangular faces; in green), and 30 edge midpoints (in red). Projecting the icosahedron from its center to a sphere maps a tessellation of the sphere into 120 triangles, as shown in Fig 2B left. The angles are π/2 for red, π/3 for green, and π/5 for blue. The generator *I* on Riemann sphere is:

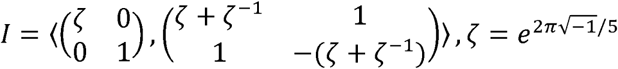

**Fig 2.**
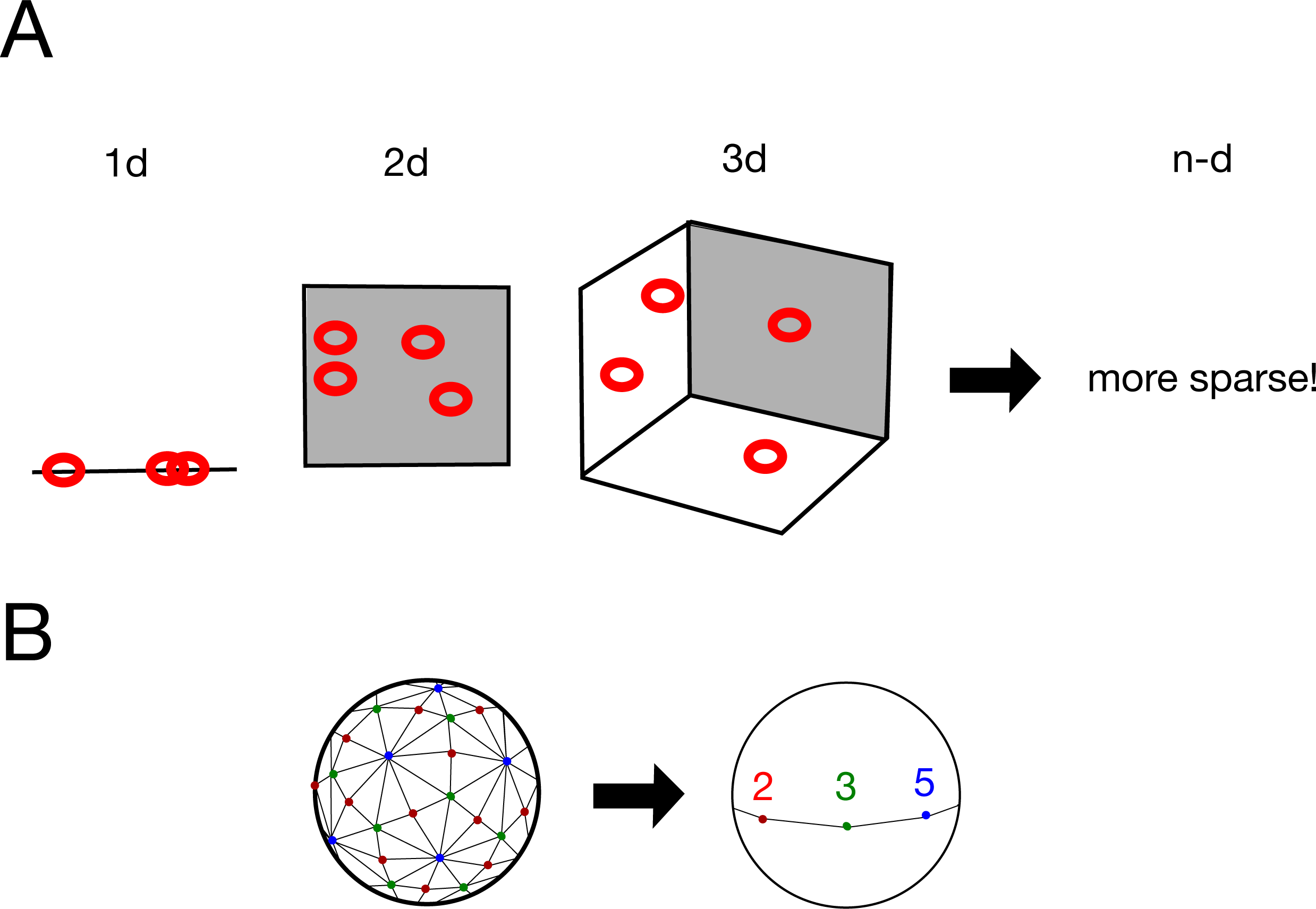
“Curse of dimensionality” and a possible solution. (A) “Curse of dimensionality” effects when there is a sparse geometric distribution of data points; see also Ronan *et al.* (2016). (B) Example of geometric conversion to a simpler system: quotient of icosahedral tessellation by *I* on a Riemann sphere. 2 (red points), 3 (green points), and 5 (blue points) correspond to the midpoints of the edges, the barycenters of the faces, and the vertices, respectively; see also Cornelissen and Kato (2005).

An icosahedron has 6 cyclic subgroups of order 5, 10 cyclic subgroups of order 3, and 15 cyclic subgroups of order 2. The quotient of this Riemann sphere by the group *I* is shown in Fig 2B right. In the figure, 2 (red points), 3 (green points), and 5 (blue points) correspond to the midpoints of the edges, the barycenters of the faces, and the vertices, respectively. As a result of this mapping, the system is simplified.

We define a *p*-adic metric based on rigid geometry, as described in the Methods section, and use this to pretreat the data before clustering or machine learning; the results are shown in Fig 1B. The control samples and those stored for long term formed distinct clusters with each of the proposed methods; this suggests this method eliminates the “curse of dimensionality”. Additionally, PCA showed 98% of contributions was attributed to PC1, PC2, PC3 and PC4. Considering absolute values of correlations, PC1 is attributed to 3y; PC2 is 2y2, 2y3; PC3 is 2y1; PC4 is f-2 with milder correlation to other control samples (Table 3). Proper clustering of p-adic values is thus well achieved also in PCA. For nMDS, the effect of clustering was prominent. All the control samples localized at almost an identical point, while samples with years of freezing environment sparse along the plotting (Fig 1B). The means of the variances in the original method and the rigid geometry method (95% confidence intervals: 60 ± 10 and 6000 ± 8000, respectively) do not reflect the advantage of using the *p*-adic metric. However, the data from the rigid method had 10 outliers in high ranks (see Fig 3A for the skewness); these were defined as being larger than the corresponding Euclidean value of the same rank. When the ten samples with the largest variances were excluded, the means of the variances obtained with the original method and the rigid geometry method became 44 ± 5 and 14 ± 3, respectively, with *p* = 5 × 10^−20^ for the *t*-test; again, this suggests the “curse of dimensionality” has been removed. We note that neural-network-based machine learning showed the same tendency as in the previous section; the number of unknown parameters was 1630. All the data indicate successful clustering with *v* metric.

**Fig 3.**
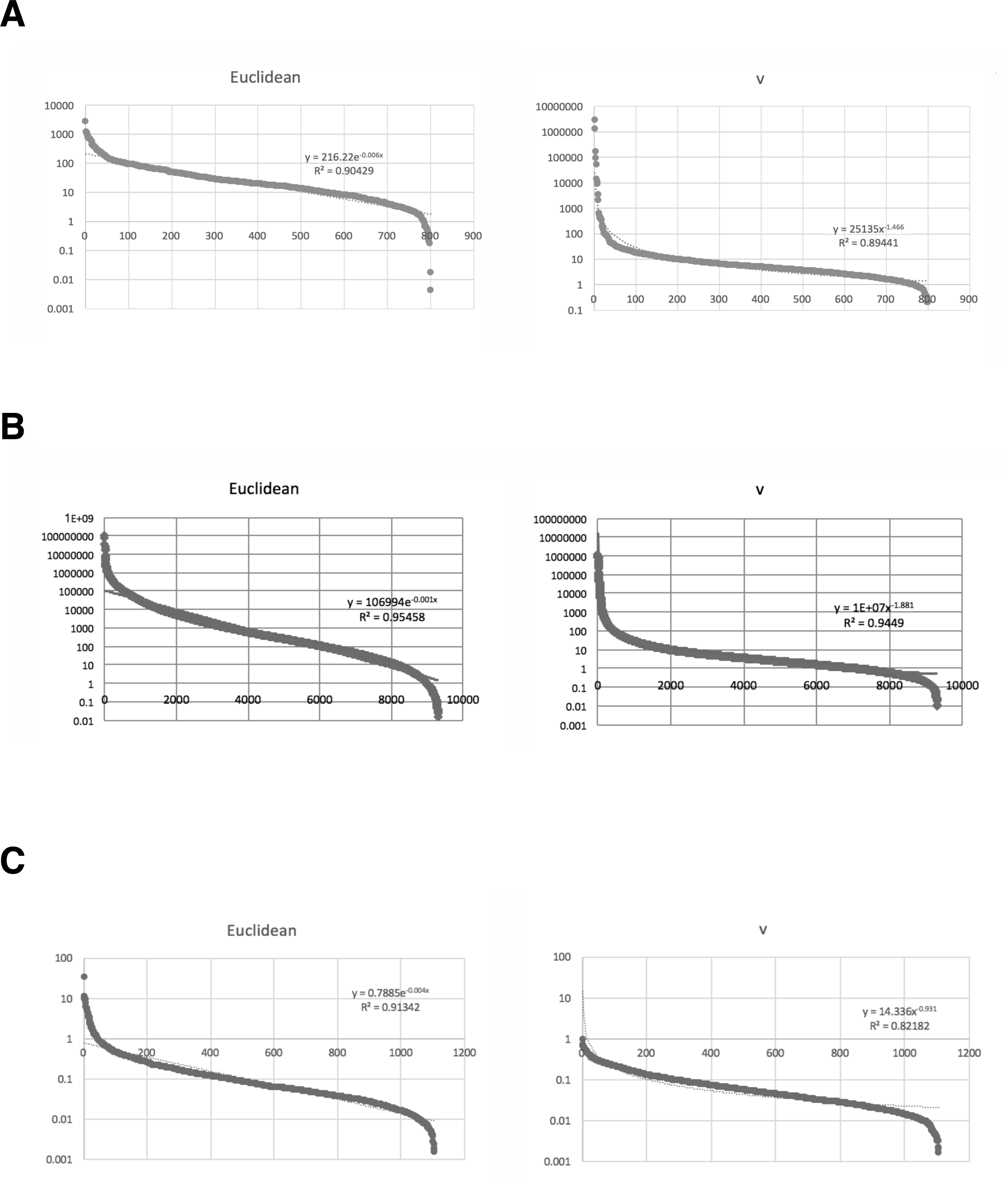
Ranked variance distributions of original values and *v* values of omics data. Euclidean; raw values. *v*; *p*(*I*)-adic *v*values. Horizontal axis: the rank of values. Vertical axis: the variances. (A) Ranked variance distributions of unused values and *v* values of proteins used for calculations (*N* = 803). (B) Ranked variance distributions of raw signal intensity values and *v* values of expression arrays for *Saccharomyces cerevisiae* used for calculations (*N* = 9335). (C) Ranked variance distributions of 2^induction factor^ values and *v*values of expression arrays for *Escherichia coli* used for calculations (*N*=1109).

**Table 3.** Correlation matrix among *v* metric of LC/MS data and principal components.

| $\nu$ | f-1 | f-2 | f-3 | 1h1 | 1h2 | o/1 | o/2 | 2y1 | 2y2 | 2y3 | 3y |
| --- | --- | --- | --- | --- | --- | --- | --- | --- | --- | --- | --- |
| PC1 | 0.14 | -0.00 | 0.06 | 0.08 | 0.19 | -0.42 | 0.23 | 0.01 | 0.01 | 0.01 | -1.00 |
| PC2 | -0.05 | 0.01 | -0.04 | 0.20 | 0.01 | -0.09 | -0.82 | 0.05 | -1.00 | 0.98 | -0.00 |
| PC3 | 0.01 | 0.03 | 0.10 | 0.05 | 0.21 | 0.08 | 0.02 | -1.00 | 0.01 | 0.06 | -0.00 |
| PC4 | -0.27 | -1.00 | -0.08 | -0.14 | -0.19 | -0.11 | -0.12 | -0.03 | -0.01 | -0.02 | 0.00 |

## Some other metrics failed to eliminate “curse of dimensionality” effects with LC/MS data

To further clarifying the outperformance of *v* metric, we tested nearest neighbor distances for the calculation of clustering in LC/MS. None of the seven hierarchical clustering methods or *k* = 5 means has successful results in clustering (Fig 4). From the original values of nearest neighbor distances, control samples and samples undergone long-term preservations were not properly clustered, either (Fig 4). Since there is only single dimension for comparison, neural network and PCA were not performed. There are also zero distances for nMDS, and this method was not performed, either.

**Fig 4.**
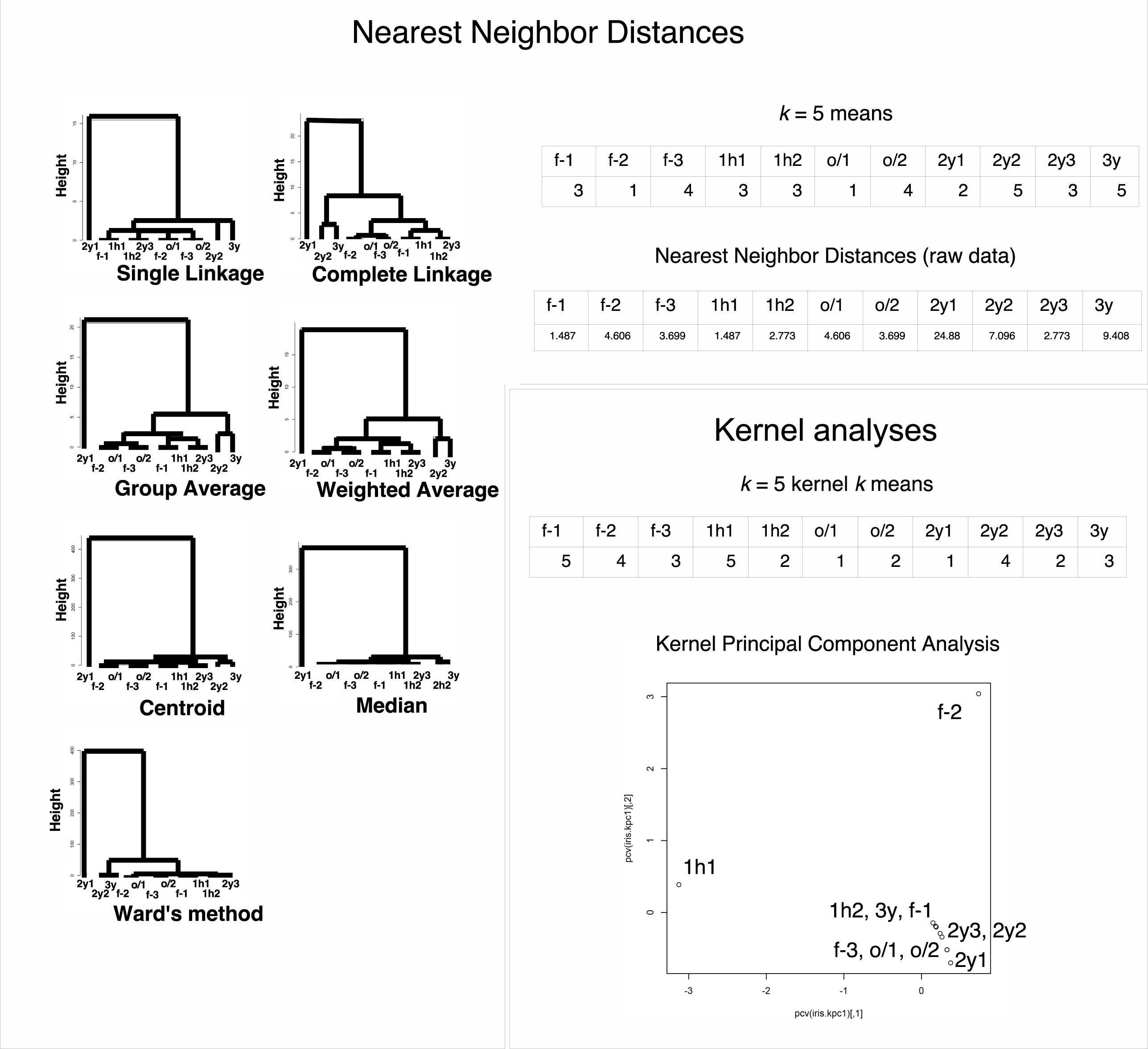
Clustering of nearest neighbor distance value sets and radial basis kernel value sets for each protein in the LC/MS data of HEK-293 (*N* = 803). f-1, f-2, and f-3: freshly prepared samples in replicates 1, 2, and 3, respectively; 1h1 and 1h2: samples frozen at _80°C for 1 h in replicates 1 and 2, respectively; o/1 and o/2: samples that remained at _80°C overnight and then in liquid nitrogen overnight (“o/n-o/n”) in replicates 1 and 2, respectively; 2y1, 2y2, and 2y3: samples preserved in liquid nitrogen storage according to the RIKEN protocol for approximately 2 years in replicates 1, 2, and 3, respectively; 3y: sample preserved in liquid nitrogen storage according to the RIKEN protocol for approximately three years. The numbers in the *k*-means table are the indices of the classified groups. The numbers in the neural network table are the cl factors, which represents the one-dimensional characteristics of the systems. The differences in cl values show dissimilarity of the samples. Please also see the Methods section.

We also performed kernel principal component analysis with radial basis kernels in LC/MS. Even utilizing this method, control samples and samples undergone long-term preservations were not properly clustered (Fig 4). For other methods such as cluster analysis via nonparametric density estimation with radial basis kernels and kernel *k*-means, neither of them had successful clustering results (Fig 4, cluster analysis via nonparametric density estimation was converged to a single group). Neural network used in this study was already with radial basis kernels, and nMDS was not suitable for the analysis because kernel is metric based.

Next, we calculated correlation distances and performed all the seven hierarchical clustering methods, *k* = 5 means, neural network, PCA and nMDS. We found that none of the method was able to achieve proper separation between long-term storage samples and frozen samples (Fig 5, Table 4). Table 4 for PCA shows PC1 and PC2 (89 and 7% contribution to the data, respectively) were the major components and PC1 lacks f-2 and 2y2 contribution, while PC2 is attributed to these two, which does not make sense for the proper clustering.

**Fig 5.**
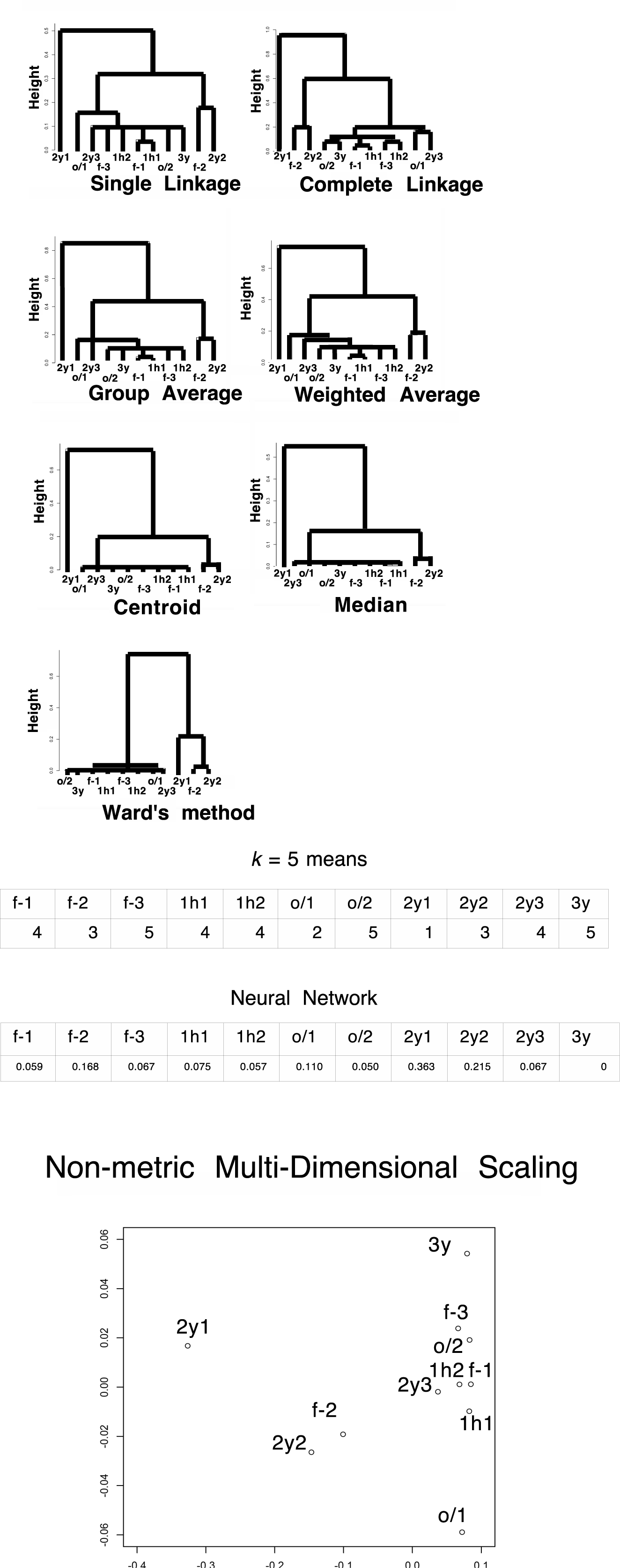
Clustering of correlation distance value sets for each protein in the LC/MS data of HEK-293 (*N* = 803). Euclidean; raw values. *v*; *p*(*I*)-adic *v* values. Horizontal axis: the rank of values. Vertical axis: the variances. (A) Ranked variance distributions of unused values and *v* values of proteins used for calculations *(N* = 803). (B) Ranked variance distributions of raw signal intensity values and *v* values of expression arrays for *Saccharomyces cerevisiae* used for calculations (N = 9335). (C) Ranked variance distributions of 2^induction factor^ values and *v* values of expression arrays for *Escherichia coli* used for calculations (*N*=1109).

**Table 4.** Correlation matrix among correlation distances of LC/MS data and principal components.

|  | f-1 | f-2 | f-3 | 1h1 | 1h2 | o/1 | o/2 | 2y1 | 2y2 | 2y3 | 3y |
| --- | --- | --- | --- | --- | --- | --- | --- | --- | --- | --- | --- |
| PC1 | 0.98 | -0.12 | 0.98 | 0.98 | 0.99 | 0.91 | 1.00 | -0.97 | -0.62 | 0.94 | 0.97 |
| PC2 | 0.11 | 0.96 | -0.02 | 0.11 | 0.10 | 0.23 | -0.03 | 0.17 | 0.76 | 0.26 | -0.03 |

## A *p*-adic metric based on rigid geometry also eliminated “curse of dimensionality” effects on some microarray data

To further characterizing the metric in other omics data such as gene expression data of microarray, we utilized existing data set of heme regulatory network in yeast *Saccharomyces cerevisiae* from [17]. In the original signal intensity data set of the expression array, heme deficient samples and heme sufficient samples were differentially clusterized in single linkage analysis and *k* = 4 means method, but failed in other six hierarchical clustering methods or neural network tested (Fig 6A). However, utilizing *v* metric, all the seven hierarchical clustering methods and *k* = 4 means method exhibited differentially clusterized heme deficient samples and heme sufficient samples (Fig 6B). In this data set, neural network with rigid geometry still did not improve the result of clustering, however. Additionally, PCA of raw signal values exhibited 97% of the data can be explained by a single component with absolute correlation values > 0.96 in each, indicating failure in clustering (Table 5). Even in *v* metric, 54% contribution was from PC1 (0.99 correlated with s2), 23% was from PC2 (-0.93 correlated with s3), 19% was from PC3 (0.96 correlated with s1), 2% was from PC4 (1.00 correlated with d3), 2% was from PC5 (1.00 correlated with d2), 0.1% was from PC6 (0.99 correlated with d1). This independency of the components shows that PCA has failed in clustering even in *v* metric (Table 6). For nMDS, original raw signal intensities showed weak clustering (Fig 6A). However, as in LC/MS data, *v* metric exhibited that all the heme-deficient samples localized at almost an identical point, while heme-sufficient samples sparse along the plotting (Fig 6B), suggesting nMDS worked very fine. The average of variances in original signals and *v* metric were 80000 ± 30000 and 400000 ± 500000, respectively (95% confidential). When we removed 39 outlier samples of *v* metric (Fig 3B), they became 70000 ± 30000 and 1400 ± 600 with *p* = 7 × 10^−6^ for the *t*-test, as in the LC/MS data of HEK-293. The number of unknown parameters in neural-network-based machine learning were 18685 and 18683 for original signals and *v* metric, respectively, indicating too many number of dimensions caused failure of clustering in neural network.

**Fig 7.**
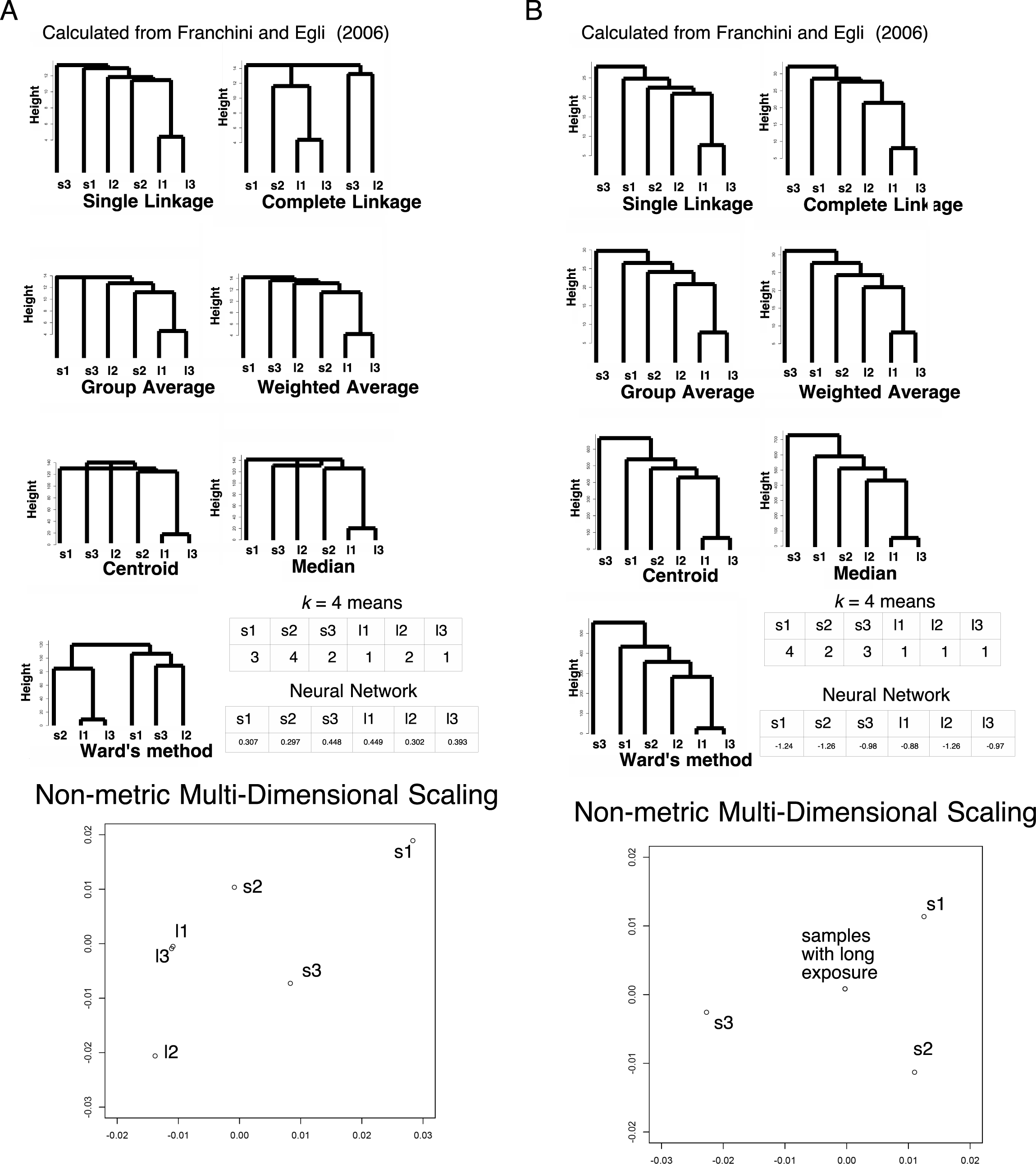
Clustering in the expression array data of *Escherichia coli (N* = 1109). s1, s2, and s3: samples with short-term starvation for glucose in replicates 1 (GSM106337), 2 (GSM106338), and 3 (GSM106339), respectively; l1, l2, and l3: samples with long-term starvation for glucose in replicates 1 (GSM106340), 2 (GSM106341), and 3 (GSM106342), respectively. The numbers in the k-means table are the indices of the classified groups. The numbers in the neural network table are the cl factors, which represents the one-dimensional characteristics of the systems. The differences in cl values show dissimilarity of the samples. Please also see [18] and the Methods section for more detail. (A) 2^induction factor^ values. (B) *v* values.

**Table 5.** Correlation matrix of microarray data in *Saccharomyces cerevisiae* and principal components. d; replicas of heme-deficient samples. s; replicas of heme-sufficient samples.

| raw | d1 | d2 | d3 | s1 | s2 | s3 |
| --- | --- | --- | --- | --- | --- | --- |
| PC1 | 0.96 | 0.99 | 0.98 | 0.99 | 0.99 | 0.99 |
| PC2 | 0.26 | 0.01 | 0.15 | -0.10 | -0.15 | -0.14 |

**Table 6.** Correlation matrix among *v* metric of microarray data in *Saccharomyces cerevisiae* and principal components. d; replicas of heme-deficient samples. s; replicas of heme-sufficient samples.

| $\nu$ | d1 | d2 | d3 | s1 | s2 | s3 |
| --- | --- | --- | --- | --- | --- | --- |
| PC1 | -0.06 | -0.02 | -0.00 | 0.01 | 0.99 | 0.30 |
| PC2 | 0.02 | -0.01 | 0.01 | -0.27 | 0.13 | -0.93 |
| PC3 | 0.05 | -0.07 | -0.04 | 0.96 | 0.03 | -0.22 |
| PC4 | 0.06 | 0.03 | 1.00 | 0.01 | 0.00 | 0.00 |
| PC5 | 0.14 | 1.00 | -0.02 | 0.01 | 0.00 | -0.00 |
| PC6 | 0.99 | -0.01 | -0.00 | -0.00 | 0.00 | 0.00 |

**Fig 6.**
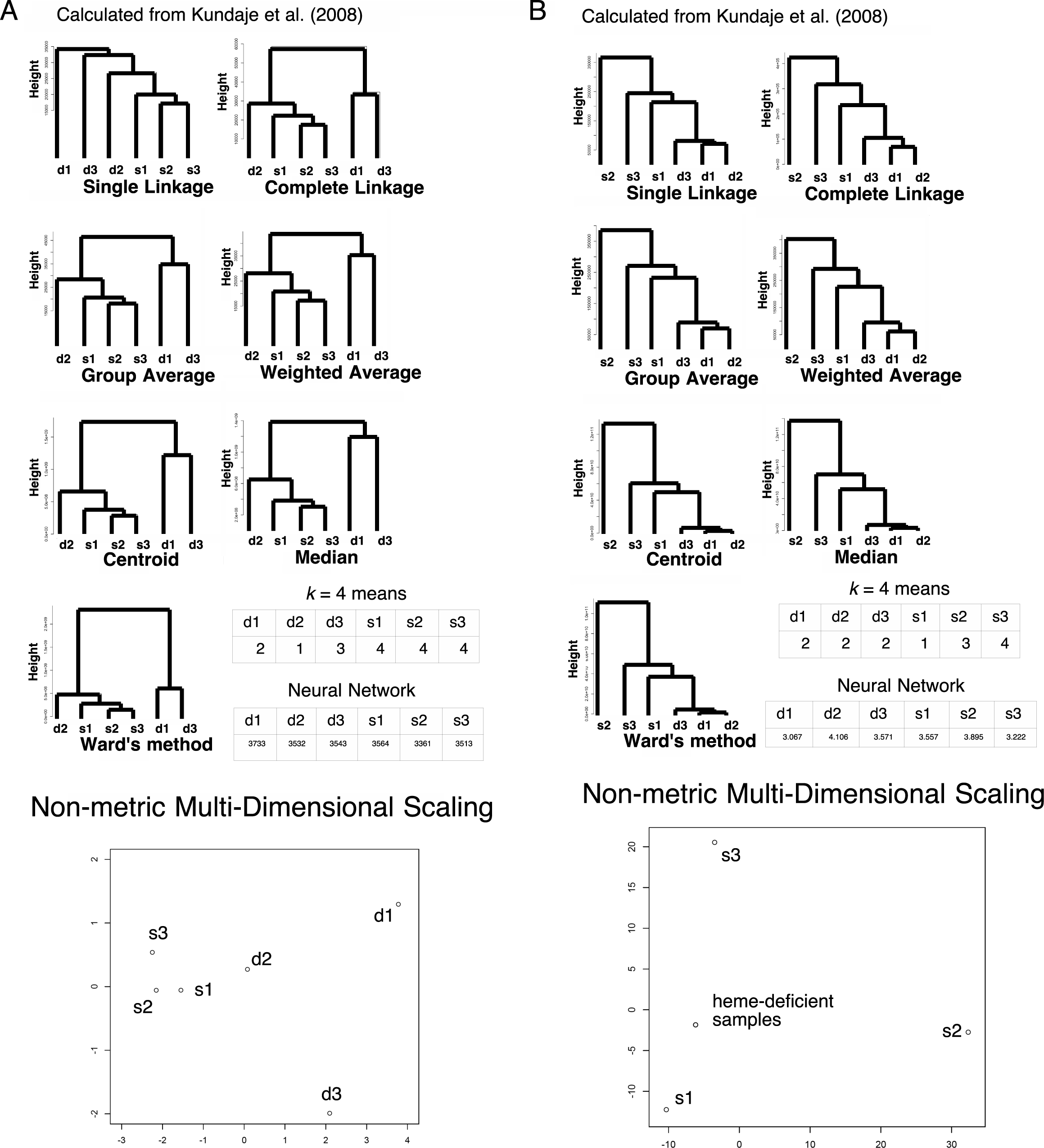
Clustering in the expression array data of *Saccharomyces cerevisiae (N* = 9335). d1, d2, and d3: heme-deficient samples in replicates 1 (GSM206793), 2 (GSM206794), and 3 (GSM206795), respectively; s1, s2, and s3: heme-sufficient samples in replicates 1 (GSM206796), 2 (GSM206797), and 3 (GSM206798), respectively. The numbers in the k-means table are the indices of the classified groups. The numbers in the neural network table are the cl factors, which represents the one-dimensional characteristics of the systems. The differences in cl values show dissimilarity of the samples. Please also see [17] and the Methods section for more detail. (A) Raw signal intensities. (B) *v* values.

We also analyzed gene expression data characterized by induction factor of expression array in the culture of *Escherichia coli* in low glucose environments from [18]. Utilizing the values calculated by 2^induction factor^, all the seven hierarchical clustering methods, *k* = 4 means and neural network failed in proper clustering between the samples of short starvation and long starvation (Fig 7A). However, the clustering of *v* metric by all the 7 hierarchical methods and *k* = 4 means has succeeded in proper arrangements of clustering (Fig 7B). For neural network, only replicate 2 of long starvation (GSM106341) was misclustered, suggesting the improvement of clustering. The number of unknown parameters in neural-network-based machine learning were 2233 and 2223 for 2^induction factor^ and *v* metric, respectively. Additionally, PCA exhibited 63% of the data can be explained by a single component with absolute correlation values > 0.71 in each. 12% of the data can be explained by s1 with a correlation value 0.54, 11% of the data can be explained by l2 with a correlation value 0.54, 7% of the data can be explained by s3 with a correlation value -0.49, and 6% of the data can be explained by s2 with a correlation value 0.46. This mild independency indicates failure in clustering (Table 7). For *v* metric, 50% of the data can be explained by a single component with absolute correlation values > 0.62 in each. 18% of the data can be explained by s3 with a correlation value 0.71, 14% of the data can be explained by s1 with a correlation value -0.74, 10% of the data can be explained by l2 with a correlation value 0.70, 7% of the data can be explained by l1 & l3 with a correlation values -0.54 and -0.53. This mild independency indicates failure in clustering (Table 8). For nMDS, original values showed weak clustering (Fig 7A). However, as in LC/MS data, *v* metric exhibited that all the samples with long exposure to a low glucose environment localized at almost an identical point, while samples with short exposure sparse along the plotting (Fig 7B), suggesting nMDS worked very fine. The average of variances in original signals and *v* metric were 0.30 ± 0.08 and 0.088 ± 0.006 (95% confidential), respectively, with *p* = 4 x 10^−7^ for the *t*-test (Fig 3C).

**Table 7.** Correlation matrix of microarray data in *Escherichia coli* and principal components. s; replicas with short-term starvation. l: replicas with long-term starvation.

| raw | s1 | s2 | s3 | l1 | l2 | l3 |
| --- | --- | --- | --- | --- | --- | --- |
| PC1 | 0.71 | 0.86 | 0.76 | 0.83 | 0.76 | 0.84 |
| PC2 | 0.54 | -0.10 | 0.26 | -0.44 | 0.09 | -0.43 |
| PC3 | -0.40 | -0.12 | 0.27 | -0.16 | 0.54 | -0.14 |
| PC4 | 0.18 | -0.15 | -0.49 | 0.04 | 0.35 | 0.06 |
| PC5 | -0.07 | 0.46 | -0.19 | -0.19 | 0.05 | -0.20 |

**Table 8.**
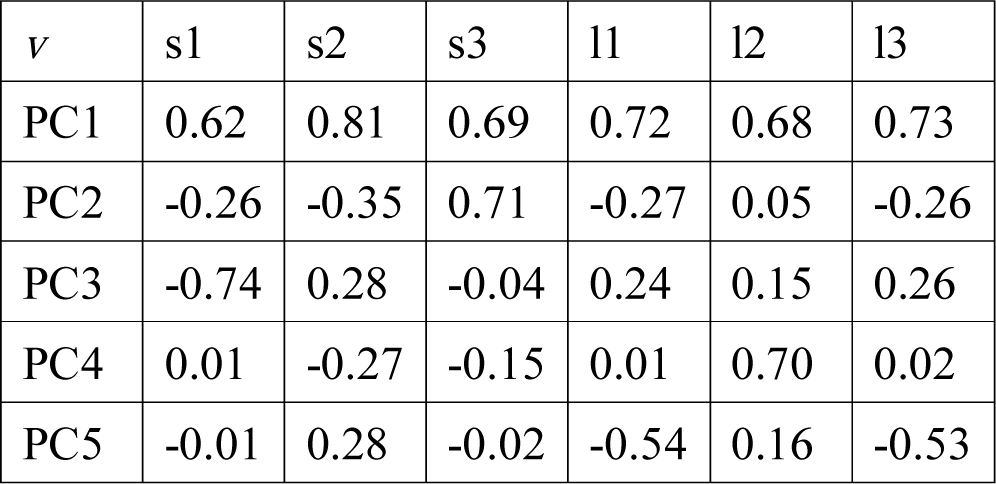
Correlation matrix among *v* metric of microarray data in *Escherichia coli* and principal components. s; replicas with short-term starvation. l: replicas with long-term starvation.

| $\nu$ | s1 | s2 | s3 | l1 | l2 | l3 |
| --- | --- | --- | --- | --- | --- | --- |
| PC1 | 0.62 | 0.81 | 0.69 | 0.72 | 0.68 | 0.73 |
| PC2 | -0.26 | -0.35 | 0.71 | -0.27 | 0.05 | -0.26 |
| PC3 | -0.74 | 0.28 | -0.04 | 0.24 | 0.15 | 0.26 |
| PC4 | 0.01 | -0.27 | -0.15 | 0.01 | 0.70 | 0.02 |
| PC5 | -0.01 | 0.28 | -0.02 | -0.54 | 0.16 | -0.53 |

These two data sets indicate that the calculation of *v* metric is effective not only in proteomics data, but also in expression data such as raw expression profiles or induction factors, at least in clustering methods and nMDS.

## Discussion

Utilizing *p*-adic rigid geometry, we succeeded in eliminating the “curse of dimensionality” effects from significantly diverged sample data, at least for the LC/MS data for HEK-293. If the total number of dimensions *(n* = Σ*nd)* exceeds the original number of model dimensions (in this case, *n* = 1630), the values converged. We assumed the number of traces Σn_*d*_–*n*) became nilpotent [19]. Here, 800 (the first rank protein data were removed from *N =* 803) × 11 = 8800 > 1630, and the observed convergence of *v* was expected beyond the underdetermined system. To support this idea, clusters f-1, f-2, and 2y2, which were misclustered in Fig 1A, were appropriately clustered when using the *v* metric (Fig 8); this was true for all the clustering methods and neural network considered, with 800 × 3 = 2400 > 1606. This allowed us to determine whether a given sample was from a nearly fresh culture or had been stored for a long time at low temperature. It is also notable that for PCA, 90, 10 and 0.4% contributions were from PC1, PC2 and PC3, and 2y2, f-2 and f-1 were correlated independently to PCs with 1.00, 1.00 and 0.97, respectively. PCA seems not to work well in high-dimensional data. For nMDS, only three points are not proper to observe clustering.

**Fig 8.**
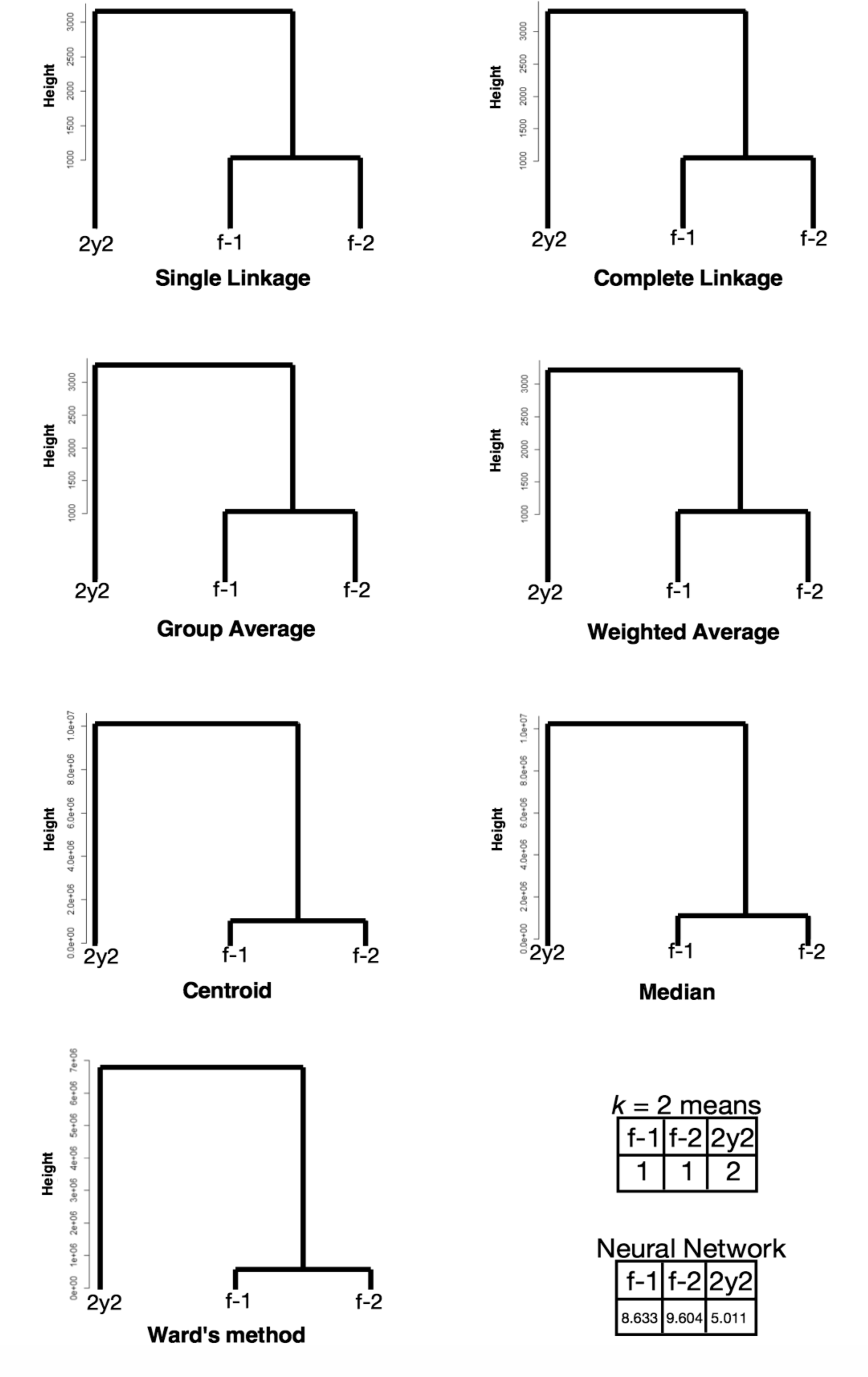
Clustering of newly invented *v* value sets of each protein in LC/MS of HEK-293 (*N* = 803). f-1 and f-2: freshly prepared samples in replicates 1 and 2, respectively; 2y2: a sample preserved in liquid nitrogen storage according to the RIKEN protocol for approximately two years in replicate 2. The numbers in the *k*-means table are the indices of the classified groups. The numbers in the neural network table are the cl factors, which represents the one-dimensional characteristics of the systems. The differences in cl values show dissimilarity of the samples. Please also see the Methods section.

This success is entirely based on an algebraic, analytic, and topological geometric analysis based on rigid geometry. So far as we know, this is the first application of “rigid geometry” to a biological system. We note that this methodology can be applied to any type of data, providing the data approximately follows a neutral logarithmic Boltzmann-type distribution. The agreement of results from a supervised machine learning and from several unsupervised clustering analyses demonstrates the power of this methodology. In biology, a similar approach can be used to evaluate the proteins inside cells of microbes [20]. The possibility of applications to other research fields, such as chemistry, physics, astronomy, and earth science, is promising but depends on successfully introducing the concept of “fitness” to the particular application.

We expect physiological differences affect the patterning of protein/mRNA expression in the type of analysis presented here. However, if fluctuations within a set of replicates are prominent, the clustering may be improperly achieved. We measure the fluctuations by observing obvious differences among control and experimental samples. From Fig 1A/Fig 6A/Fig 7A, obviously all the hierarchical clustering analyses show contradictory results, while from Fig 1B/Fig 6B/Fig 7B, all the hierarchical clustering analyses show matched results, indicating significant improvement at least for hierarchical clustering. Control samples of fresh samples vs. very short time of freezing have statistically non-significant difference in average quantification (*p* = 0.40 for *t*-test), while that vs. years of freezing do have statistically significant difference (*p* = 6.0 x 10^−8^ for *t*-test). We thus concluded that long-term storage in low temperature does have affect sample preservation, while short-term does not.

We tried to utilize neural network for an example of supervised approach, which is more powerful tool for classification. However, the approach was not as trustable as we had expected, according to incalculable Akaike information criterion. Furthermore, the data of Fig 6A and Fig 6B have suggested neural network is not trustable if the data have very high dimensions. We thus moved to more general approach of clustering that is trustable in any methods we have tested, and this is the aim of this work. Our approach seems to erase the traces of huge degree of freedom by erasing the singularities. Additionally, regression analysis still depends heavily on the number of independent variables. If the number of variables is still large, it is difficult to classify the data due to “curse of dimensionality”. The data of PCA and *p*-adic metric performed well in LC/MS compared with failure of clustering in original unused vales. However, neither original values nor *v* metrics perform well in expression arrays. This might be that high dimensionality prevents reducing dimensions to certain small values in PCA. It is also notable that in nMDS, control samples clustered to almost a single point, while experimental samples were sparse along the plots, indicating clear distinction of the characteristics of the samples by nMDS.

In conclusion, we have succeeded in removing the “curse of dimensionality” from the observed differences among control and treatment (fresh and stored) samples of HEK-293 cells when evaluating LC/MS data, and other expression data of *Saccharomyces cerevisiae* and *Escherichia coli* treated by clustering methods and nMDS. The success was entirely based on the topological characteristics of the *p*-adic metric on rigid geometry. This approach has the potential to calculate the characteristic values of any system for which the data approximate a neutral logarithmic Boltzmann distribution.

## Materials and Methods

### Cell culture

A human HEK-293 cell line from an embryonic kidney was purchased from RIKEN (Japan). The original cultures were frozen on either March 18, 2013 (3-year storage) or March 5, 2014 (2-year storage), and they were used in experiments between February and June 2016. The strain was cultured in Modified Eagle’s Medium (MEM) + 10% fatal bovine serum (FBS) + 0.1 mM nonessential amino acid (NEAA) at 37°C with 5% CO2. Subculturing was performed in 0.25% trypsin, and prior to the experiment, the original cells from RIKEN were frozen following the standard protocol provided by RIKEN: in culture medium with 10% dimethyl sulfoxide (DMSO), they were cooled until reaching 4°C at –2°C/min, held at that temperature for 10 min, cooled until reaching –30°C at –1°C/min in order to freeze, held at that temperature for 10 min, cooled again until reaching –80°C at –5°C/min, and then held at that temperature overnight. The next day, they were transferred to storage in liquid nitrogen. Freezing conditions for the control samples are described in the Results section.

### Protein extraction, alkylation, and digestion

The HEK-293 proteins were extracted using the standard protocol for the RIPA buffer (NACALAI TESQUE, INC., Kyoto, Japan). Approximately 10^6^ harvested cells were washed once in Krebs-Ringer-Buffer (KRB; 154 mM NaCl, 5.6 mM KCl, 5.5 mM glucose, 20.1 mM HEPES pH 7.4, 25 mM NaHCO_3_). They were resuspended in 30 µl of RIPA buffer, passed in and out through 21G needles for destruction, and incubated on ice for 1 h. They were then centrifuged at 10,000 *g* for 10 min at 4°C, followed by collection of the supernatants; the proteins were quantified by using a Micro BCA Protein Assay Kit (Thermo Fisher Scientific, Waltham, U.S.A.), and further processing was performed using XL-Tryp Kit Direct Digestion (APRO SCIENCE, Naruto, Japan). The samples were solidified in acrylamide gel, washed twice in ultrapure water, washed three times in dehydration solution, and dried. The samples were then processed using an In-Gel R-CAM Kit (APRO SCIENCE, Naruto, Japan). The samples were reduced for 2 h at 37°C, alkylated for 30 min at room temperature, washed five times with ultrapure water, washed twice with destaining solution, and then dried. The resultant samples were trypsinized overnight at 35°C. The next day, the dissolved digested peptides were collected by ZipTipC18 (Merck Millipore, Corp., Billerica, U.S.A.). The tips were dampened with acetonitrile twice and equilibrated twice with 0.1% trifluoroacetic acid. The peptides were collected by ~20 cycles of aspiration and dispensing, washed twice with 0.1% trifluoroacetic acid, and eluted by 0.1% trifluoroacetic acid /50% acetonitrile with aspiration and dispensing five times × three tips followed by vacuum drying. The finalized samples were stored at -20°C. Before performing LC/MS, they were resuspended in 0.1% formic acid, and the amounts were quantified by Pierce Quantitative Colorimetric Peptide Assay (Thermo Fisher Scientific, Waltham, U.S.A.). This protocol is published at http://dx.doi.org/10.17504/protocols.io.h4qb8vw.

### LC/MS

LC/MS was performed by the Medical Research Support Center, Graduate School of Medicine, Kyoto University with a quadrupole–time-of-flight [Q-Tof] mass spectrometer TripleTOF 5600 (AB Sciex Pte., Ltd., Concord, Canada). We followed their standard protocols. The loading amount for each sample was 1 µg. We extracted the quantitative data for the unused information for identified proteins by using ProteinPilot 4.5.0.0 software (AB Sciex Pte., Ltd., Concord, Canada).

### Clustering analyses and machine learning of the pattern

For LC/MS, hierarchical clustering analyses were performed by the standard hclust function in R 3.2.3 (https://cran.r-project.org) with the package stats. The actual hierarchical methods used were: single linkage, complete linkage, group average, weighted average, centroid, median, and Ward’s method. The *k*-means method was performed by the standard kmeans function in R 3.2.3 with the package stats. It was calculated based on all eleven/six samples. For machine learning, an 11-1-1 hierarchical neural-network analysis was performed in R 3.2.3 with the package nnet, and the cl (number of raw data points) factors were calculated as a characterization index for the pattern. For expression arrays, all the analyses were done the same as LC/MS except using R 3.3.2 and 6-1-1 hierarchical neural-network. Principal component analyses (PCA) and non-metric multi-dimensional scaling (nMDS) were also performed by prcomp and isoMDS functions of R 3.3.2 with the packages stats and MASS, respectively. The metric used for nMDS was: 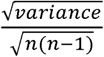. Nearest neighbor distances were calculated by nndist function of R 3.3.2 with the package spatstat. Kernel principal component analysis was performed by kpca function of R 3.3.2 with the package kernlab, selecting rbfdot option. Cluster analysis via nonparametric density estimation was performed by pdfCluster function of R 3.3.2 with the package pdfClsuter. Kernel k-means method was performed by kkmeans function of R 3.3.2 with the package kernlab. Correlation distances were calculated by corDist function of R 3.3.2 with the package MKmisc. With correlation distances, 10-1-1 neural network was used because one of the 11 values was zero. We only used the unused values or expression data that are observable in each of the eleven/six samples; this was done to avoid distortion of the calculation from identification failures within the LC/MS process or microarray experiments, since there were relatively few signal values (*N* = 803, 9335, 1109 in each sample). The actual unused values or expression data used in the calculation are shown in S2 Table.

## Supporting Information

S1 Appendix. Utilizing a *p*-adic metric embedded in rigid geometry.

Appendix indicating calculation of the new metric *v*

S2 Table. The table of raw values for the identified proteins and expression arrays.

Sheet 1, unused values for LC/MS in HEK-293; Sheet 2, raw signal intensities for expression array in *Saccharomyces cerevisiae* [17]; Sheet 3, raw induction factor values of *Escherichia coli* in [18]. Please also see the legend of Fig 1A, 7 and 9.

## Acknowledgments

We thank Prof. Tatsuaki Tsuruyama and the Medical Research Support Center, Graduate School of Medicine, Kyoto University for performing LC/MS. Additional support was provided by the Center for Anatomical, Pathological, and Forensic Medical Research, Graduate School of Medicine, Kyoto University, Japan. We are grateful for this laboratory support.

## Author Contributions

Shun Adachi is the sole author of this article.

